# Repurposing the dark genome. IV – noncoding proteins

**DOI:** 10.1101/2023.06.29.547021

**Authors:** Sarangdhar Nayak, Pawan K. Dhar

## Abstract

The dark genome comprising of non-expressing, non-translating, and extinct DNA sequences has remained a largely unexplored genomic space. Using computational and experimental approaches, novel insights into the dark matter genome have recently been gained, revealing the presence of a vast and unexplored resource. Non-coding RNA (ncRNA) refers to a class of RNA molecules that do not encode proteins but play important regulatory roles in the cell. We asked if it was possible to make functional peptides and proteins from ncRNA leading to a new biological insight and applications? Here we present initial computational data in support of making functional noncoding proteins (NCP) from ncRNA sequences. Different types of non-coding genomic sequences originating from *Caenorhabditis elegans, Drosophila melanogaster, Arabidopsis thaliana*, and *Homo sapiens* were studied to understand sequence composition, secondary structure, and physiochemical properties of NCPs. This work builds the foundation for experimentally characterizing the first-in-the-class non-coding proteins leading to a novel insights and applications.

## 1. Introduction

The term genome refers to the complete set of genetic material or DNA present in an organism. From the functional standpoint, DNA sequences have been categorized into: sequences encoding proteins, sequences encoding RNA, and sequences that do not express. The non-expressing sequences, non-translating, and extinct DNA sequences that can be artificially encoded into functional molecules, are collectively referred as the dark genome. The non-expressing sequences may include antisense, reverse coding, repetitive sequences, intergenic sequences. The non-translating sequences may include tRNA, ncRNA, ribosomal RNA, and introns. The extinct sequences include pseudogenes that were functional in the past but were retired from expression during evolution. Our work focuses on discovering the function and purpose of these dark genome sequences.

With the increase in biological complexity, genomic proportion of protein-coding DNA has been found to generally decrease during evolution. For example, microbes dedicate approx. 90% of their genome towards encoding proteins, while in yeast it is 68%, nematodes 23-24%, and humans ∼1.5% (Shabalina and Spiridonov 2004).

In contrast to protein-coding sequences, non-coding RNA (ncRNA) genome defines a group of DNA sequences that encode RNA only. Based on the size and function, non-coding RNAs have been broadly categorized into (i) transfer RNA (tRNA): tRNA molecules that act as adapters, matching specific amino acids to their corresponding codons on messenger RNA (mRNA) during translation (ii) ribosomal RNA (rRNA), a major component of ribosomes that helps in the assembly of amino acids into protein chains based on the information carried by mRNA (iii) microRNA (miRNA) that are short RNA molecules that regulate gene expression by binding to target mRNA molecules (iv) long non-coding RNA (lncRNA) molecules that act as molecular scaffolds, guides for protein complexes, or regulators of gene expression.

In addition, there are small nuclear RNA (snRNA) involved in the splicing of pre-messenger RNA (pre-mRNA), small nucleolar RNA (snoRNA) that mostly guide chemical modifications of ribosomal RNA (rRNA) and piwi-interacting RNA (piRNA) that are involved in regulating transposable elements and maintaining genome stability.

The non-coding RNA sequences that are greater than 200 nucleotides are generally classified as long non-coding RNA. ncRNA lack the ability to encode proteins. The ncRNA biology finds its footprints in molecular events like chromatin-modifying complexes, participation in shaping nuclear domains, transcriptional enhancers and so on (Engreitz et al., 2016, Rinn et al., 2012, Bierhoff et al., 2014, Marchess et al., 2014). Additionally, some ncRNAs have been observed to disrupt the transcriptional machinery or contribute to the maintenance of nuclear speckle structures. (Prasanth et al., 2005, Clemson et al., 2009, Sunwoo et al., 2009). ncRNAs have also been found to be active regulators of splicing, mRNA decay, protein translation, protein stability, or as molecular decoys for miRNAs (Yoon et al., 2013, Quinn et al., 2016).

While the ncRNA biology has exploded in terms of scientific contributions and is an established field, the ability to synthetically produce functional peptides and proteins from ncRNA is still an unchartered territory. The idea of making functional proteins from the naturally non-expressing sequences was proposed 15 y back (Dhar et al., 2009) when growth inhibitory proteins were made by synthetically expressing E. coli intergenic sequences. After this proof-of-the-concept paper, therapeutic peptides were predicted from intergenic sequences of yeast. (Joshi et al., 2013), functional proteins were envisaged from pseudogenes (Shidhi et al., 2015). Likewise, antimicrobials, anti-Alzheimer and anti-leishmania peptides were predicted from the intergenic sequences of different model organisms (Chakrabarty et al., 2022, Deepthi et al., 2016, Raj et al., 2015).

The work took an interest turn when tRNA-encoded peptides were experimentally produced and found to have strong anti-Leishmania property (Chakrabarti et al 2022). More recently, functional protein were predicted from the antisense strand and reverse strands of DNA (Garg et al., 2023, Nayak et al., 2023).

Here, we provide computational evidence in favor of making non-coding proteins from ncRNA molecules by using genomic information from *Caenorhabditis elegans, Drosophila melanogaster, Arabidopsis thaliana*, and *Homo sapiens*.

## 2. Material and Methods

The *Caenorhabditis elegans, Drosophila melanogaster, Arabidopsis thaliana*, and *Homo sapiens* genome database was considered in this study (NCBI Resource Coordinators, 2013, LNCipedia 2013, RNAcentral 2017). The long non-coding RNA, miRNA, snRNA and snoRNA gene sequences were used in this study.

### Translation of ncRNA gene sequence

UGENE-Integrated Bioinformatics Tool was used to computationally translate noncoding RNA sequences into proteins (Okonechnikov et al., 2012). To avoid an enormous combinatorial issue, only full-length translated sequences without any stop codons were considered in this study. The Expasy-Translation Tool (Gasteiger, 2003) was used to ensure the quality of the results.

### Protein Sequence Similarity

To assess similarity, the full-length non-coding protein sequences were BLASTed against the NCBI Non-Redundant protein database. Only those sequences that showed non-similarity to existing sequences were considered. A total of 10 novel predicted proteins from each model organism were taken for detailed study (Table 1).

**Table 1:** A sample of RNA(lncRNA, snoRNA, snRNA, miRNA) proteins in *C. elegans, A. thaliana, D. melanogaster & H. sapiens* used in this study.

| S.No. | Organism | Type of ncRNA | Non-coding RNA Genes | Full-length non-coding RNA Protein |
| --- | --- | --- | --- | --- |
| 1. | *C. elegance* | lncRNA | 3154 | 118 |
|  |  | snoRNA | 346 | 46 |
|  |  | snRNA | 94 | 19 |
|  |  | miRNA | 709 | 361 |
| 2. | *D. melanogaster* | lncRNA | 42848 | 177 |
|  |  | snoRNA | 32 | 29 |
|  |  | snRNA | 27 | 0 |
|  |  | miRNA | 258 | 72 |
| 3. | *A. thaliana* | lncRNA | 4046 | 34 |
|  |  | snoRNA | 71 | 0 |
|  |  | snRNA | 66 | 13 |
|  |  | miRNA | 313 | 19 |
| 4. | *H. sapiens* | lncRNA | 55028 | 137 |
|  |  | snoRNA | 933 | 98 |
|  |  | snRNA | 1855 | 270 |
|  |  | miRNA | 1826 | 658 |

### mRNA secondary structure

The Mfold server (Zuker, 2003) was employed to calculate the mRNA secondary structure and determine its associated free energy. By combining the free energy values from different secondary structural components, an estimated total free energy for the molecule’s structure was obtained. The Mfold server performs this calculation and compares it with the relative thermodynamics of various structures derived from the same sequence (Table 3).

### Physicochemical properties

The Expasy Protparam tool, developed by the SIB Swiss Institute of Bioinformatics (Roy et al., 2011), was used to calculate various physiochemical properties such as instability index, aliphatic index, theoretical Pi, grand average of hydropathicity (GRAVY) index, and molecular weight (Table 4).

### Structure, Function Prediction, Subcellular localization

To predict the tertiary structure and potential function of proteins, the I-TASSER Server (Iterative Threading ASSEmbly Refinement) tool was employed (Zhang, 2008). The I-TASSER tool is a standard platform for its accurate predictions and generation of 3D protein structures. For determining subcellular localization, the WoLF PSORT tool was used (Horton et al., 2007). The Ramachandran plot was used to assess the distribution of amino acids in a two-dimensional space, achieved by calculating the torsional angles of amino acids φ (phi) and ψ (psi) within a protein sequence. The Procheck server was used to investigate the stereochemical properties (Table 2 & 5).

**Table 2.**
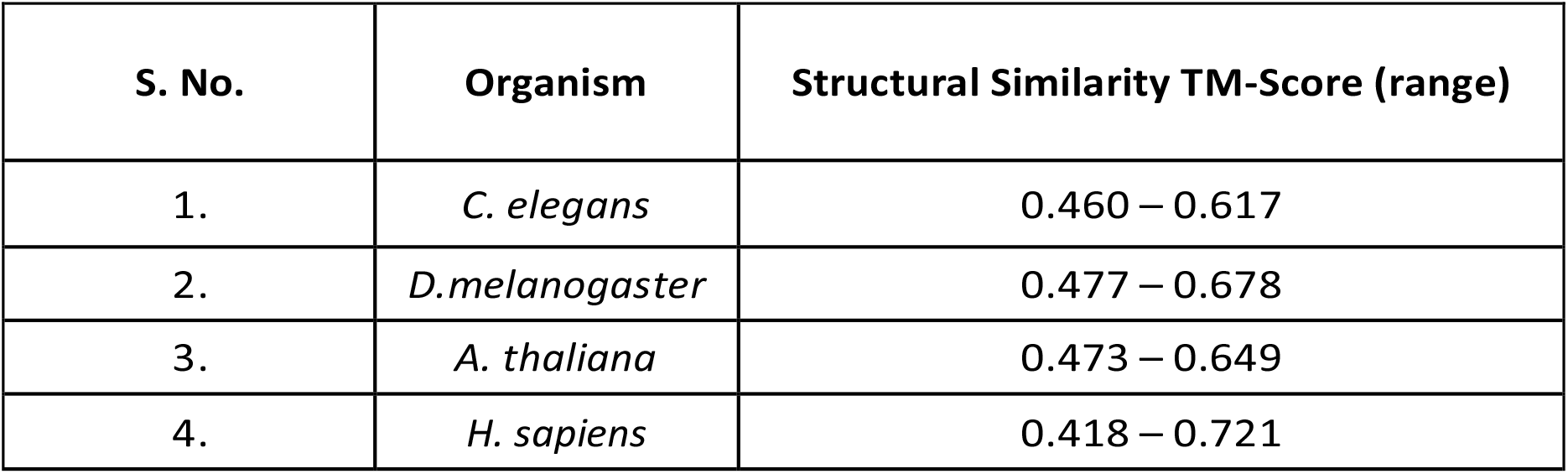
Predicted structural similarity of unique ncRNA (lncRNA, snoRNA, snRNA, miRNA) proteins of *C. elegans, D. melanogaster, A. thaliana & H. sapiens*.

## 3. Results

### Translation of ncRNA gene Sequences

On translating the ncRNA sequence of all four model organisms *C. elegans, D. melanogaster, A. thaliana*, and *H. sapiens* many full-length protein sequences were obtained without showing intervening stop codons (Table 1).

### Sequence and Structure Similarity

The full-length non coding protein sequences from *C. elegans, D. melanogaster, A. thaliana*, and *H. sapiens* were examined for their uniqueness against the existing NCBI protein sequence database (Table 1). In the case of *D. melanogaster* snRNA and *A. Thaliana* snoRNA no novel protein was found.in the rest, recently obtained novel protein sequences are completely different from the established repertoire of known protein sequences but their structure showed a strong similarity with existing proteins. The TM score above 0.5 gives high structural similarity (Table 2).

### mRNA secondary structure

The RNA molecule exhibiting the most negative free energy value is regarded as the most structurally organized and stable which mainly depends upon its length, nucleotide content and order. Interestingly, all the ncRNA proteins examined in the study were found to fall within the favorable range of free energy values, indicating their propensity for stable and well-structured secondary structures (Table 3)

**Table 3.** mRNA secondary structure free energy of putative ncRNA (lncRNA, snoRNA, snRNA, miRNA) proteins of *C. elegans, D. melanogaster, A. thaliana & H. sapiens*.

| S. No. | Organisms | mRNA secondary structure free energy (Kcal/mol) (range) |
| --- | --- | --- |
| 1. | *C. elegans* | -45.9 to -90.6 |
| 2. | *D. melanogaster* | -44.3 to -67.1 |
| 3. | *A. thaliana* | -24.5 to -65.6 |
| 4. | *H. sapiens* | -165.80 to -239.3 |

### Physicochemical properties

By analyzing the physicochemical properties of ncRNA proteins, as presented in Table 4, it was found that if these proteins are expressed within a cell, they exhibit structural stability.

**Table 4.**
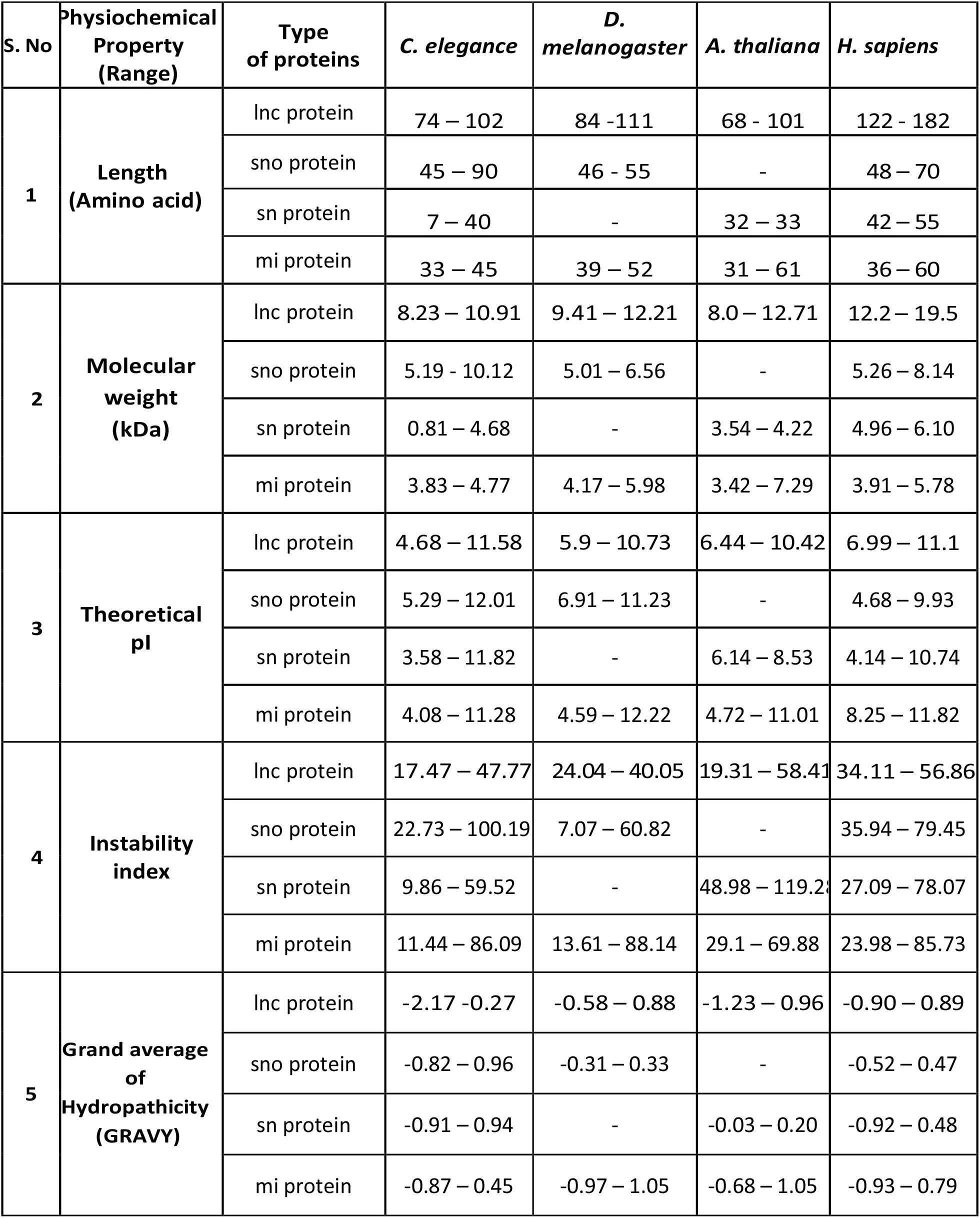
Physicochemical properties of putative ncRNA (lncRNA, snoRNA, snRNA, miRNA) proteins of C. elegans, D. melanogaster, A. thaliana & H. sapiens.

| S. No | Physiochemical Property (Range) | Type of proteins | *C. elegans* | *D. melanogaster* | *A. thaliana* | *H. sapiens* |
| --- | --- | --- | --- | --- | --- | --- |
| 1 | Length (Amino acid) | lnc protein | 74 – 102 | 84 -111 | 68 - 101 | 122 - 182 |
|  |  | sno protein | 45 – 90 | 46 - 55 | - | 48 – 70 |
|  |  | sn protein | 7 – 40 | - | 32 – 33 | 42 – 55 |
|  |  | mi protein | 33 – 45 | 39 – 52 | 31 – 61 | 36 – 60 |
| 2 | Molecular weight (kDa) | lnc protein | 8.23 – 10.91 | 9.41 – 12.21 | 8.0 – 12.71 | 12.2 – 19.5 |
|  |  | sno protein | 5.19 - 10.12 | 5.01 – 6.56 | - | 5.26 – 8.14 |
|  |  | sn protein | 0.81 – 4.68 | - | 3.54 – 4.22 | 4.96 – 6.10 |
|  |  | mi protein | 3.83 – 4.77 | 4.17 – 5.98 | 3.42 – 7.29 | 3.91 – 5.78 |
| 3 | Theoretical pI | lnc protein | 4.68 – 11.58 | 5.9 – 10.73 | 6.44 – 10.42 | 6.99 – 11.1 |
|  |  | sno protein | 5.29 – 12.01 | 6.91 – 11.23 | - | 4.68 – 9.93 |
|  |  | sn protein | 3.58 – 11.82 | - | 6.14 – 8.53 | 4.14 – 10.74 |
|  |  | mi protein | 4.08 – 11.28 | 4.59 – 12.22 | 4.72 – 11.01 | 8.25 – 11.82 |
| 4 | Instability index | lnc protein | 17.47 – 47.77 | 24.04 – 40.05 | 19.31 – 58.41 | 34.11 – 56.86 |
|  |  | sno protein | 22.73 – 100.19 | 7.07 – 60.82 | - | 35.94 – 79.45 |
|  |  | sn protein | 9.86 – 59.52 | - | 48.98 – 119.28 | 27.09 – 78.07 |
|  |  | mi protein | 11.44 – 86.09 | 13.61 – 88.14 | 29.1 – 69.88 | 23.98 – 85.73 |
| 5 | Grand average of Hydropathicity (GRAVY) | lnc protein | -2.17 -0.27 | -0.58 – 0.88 | -1.23 – 0.96 | -0.90 – 0.89 |
|  |  | sno protein | -0.82 – 0.96 | -0.31 – 0.33 | - | -0.52 – 0.47 |
|  |  | sn protein | -0.91 – 0.94 | - | -0.03 – 0.20 | -0.92 – 0.48 |
|  |  | mi protein | -0.87 – 0.45 | -0.97 – 1.05 | -0.68 – 1.05 | -0.93 – 0.79 |

These proteins possess all the characteristics in the favorable range of isoelectric point, instability index, and hydropathicity value that enable them to maintain their stability when expressed in a cellular environment.

### Subcellular Localization tertiary structure and Function Prediction

The localization of proteins provides valuable insights into their potential roles within the cell. Localization predicted by WoLF PSORT showed most proteins in cytoplasm and nucleus in worms, flies, humans and thylakoids of chloroplast in *A. thaliana*. The I-TASSER Server was employed to predict the tertiary structure of the peptide, and the corresponding C-scores for the model structures are provided in Table 5. The C-score range for the peptides was documented to span from -5 to -2.03. The function predicted based on their structure showed various enzymatic roles if they were expressed in the cell.

**Table 5.**
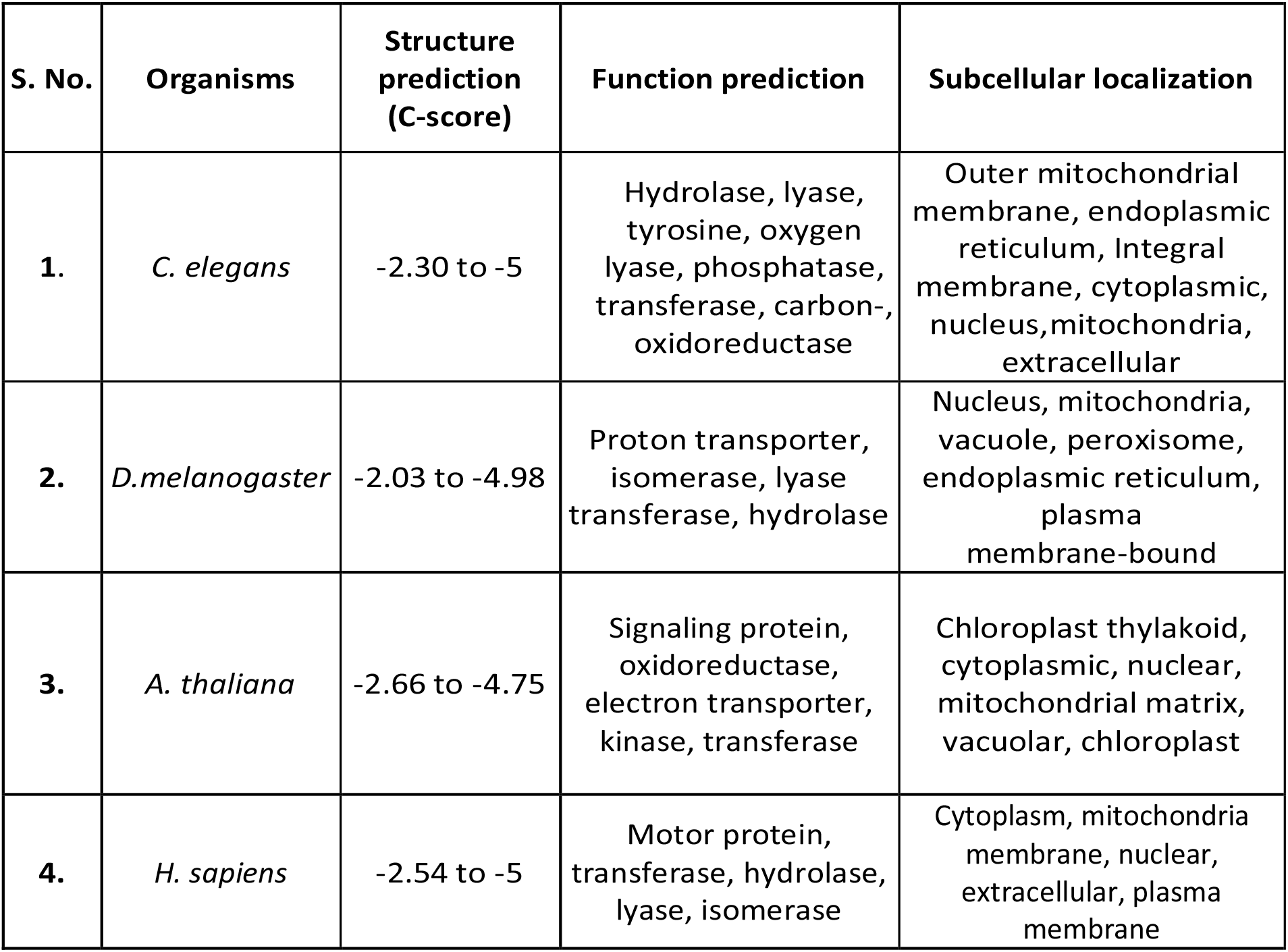
Predicted structure and function prediction of putative ncRNA (lncRNA, snoRNA, snRNA, miRNA) proteins of *C. elegans, D. melanogaster, A. thaliana & H. sapiens*.

### Stereochemical properties

For the stereochemical property, the Ramachandran plot analysis revealed a highly favorable distribution of residues (fig 2). The majority of the residues were found to occupy the preferred regions of the plot.

**Fig. 1.**
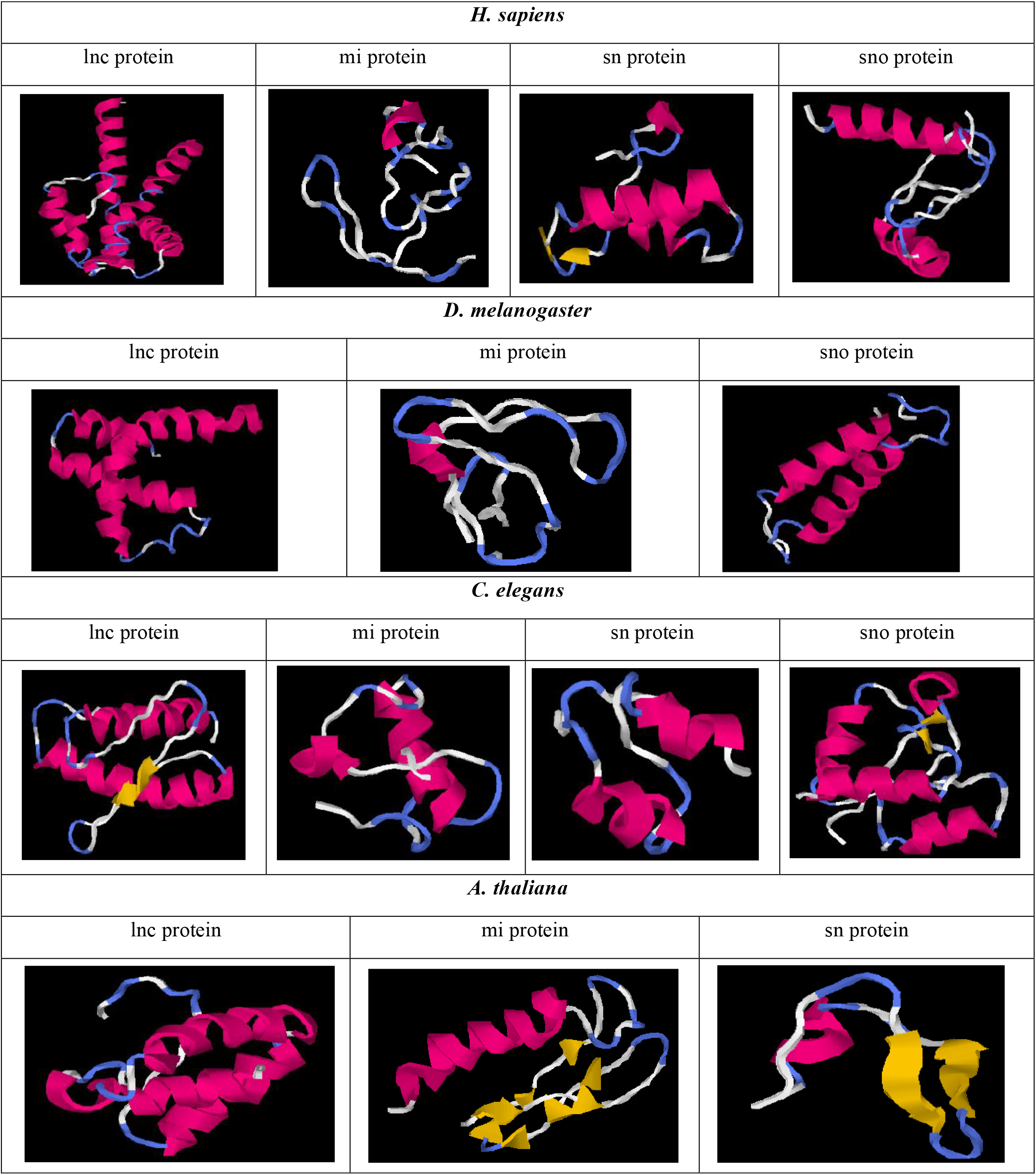
Tertiary structure of ncRNA protein resembling naturally expressing proteins (prediction tool: I-TASSER)

**Fig. 2.**
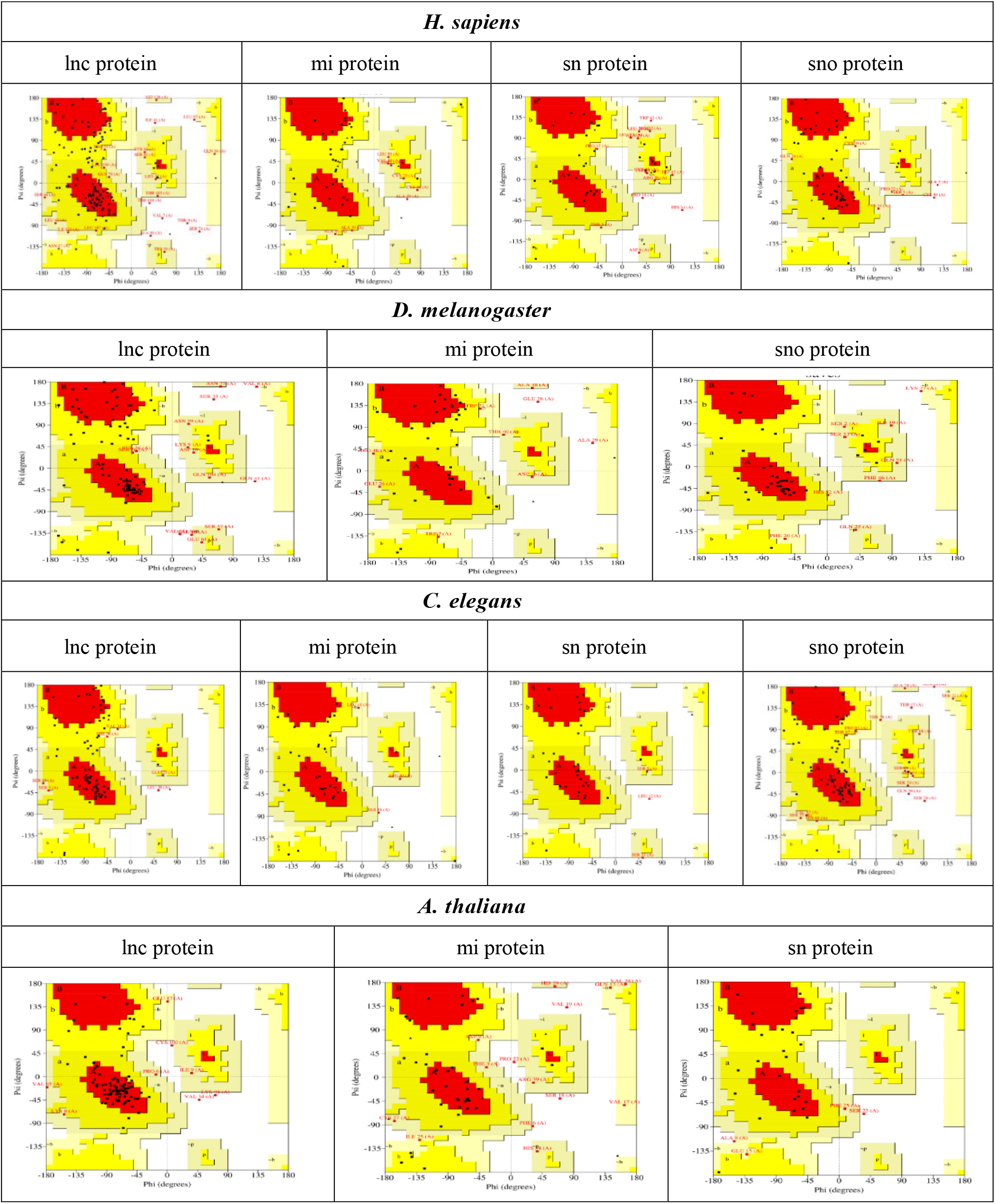
Ramachandran Plot of ncRNA proteins of C. elegans, D. melanogaster, A. thaliana, and H. sapiens.

## 4. Discussion

DNA encodes information necessary for the synthesis of proteins, the workhorses of cellular processes. To regulate the RNA and protein synthesis, bulk of the genome transcribes into non-coding RNA (ncRNA). The non-coding regions, previously thought to be functionless called “junk DNA,” have been found to play key role in gene regulation, chromosome dynamics, and other regulatory functions. It is important to note that evolution does not necessarily eliminate non-coding elements if they do not show detrimental effects on survival and reproduction. Due to the intricate and interconnected nature of gene regulation and cellular processes non-coding RNA may have been evolutionarily conserved to ensure cellular survival.

Given that cells do not graduate ncRNA sequences to encode polypeptides, we asked what if ncRNA came with an evolutionary possibility of making proteins in some organisms. How would these proteins looks like, in terms of structure, interaction and function? Finding natural evidences of non-coding proteins made from non-coding RNA would challenge our current understanding and open up exciting avenues of research.

This study was aimed at analyzing sequence and structural characteristics of proteins and along with their potential functionalities. The protein sequence equivalents were derived from the non-coding RNA and structure predicted using I-TASSER. Predictions showed up several potential enzymatic functions as hydrolase, lyase, tyrosine phosphatase, transferase, carbon-oxygen lyase, oxidoreductase, isomerase, oxidoreductase, kinase. Some of the proteins were predicted to function as proton transporter, electron transporter, signaling proteins, motor protein and so on.

Given that a cellular address determines he function and boundary condition of protein activity, most of the non-coding proteins were predicted to be localized in the cytoplasm, nucleus, and membrane. Furthermore, it was observed that most of the proteins exhibited favorable biochemical characteristics e.g., isoelectric point, hydropathy index, and instability index. These findings indicate that the noncoding proteins may possess properties that are conducive to their stability and proper folding within the cellular environment, if synthesized artificially.

The analysis of the stereochemical property by Ramachandran plot showed the distribution of residues in the preferred region, indicating potential for protein folding and stable tertiary structure, critical for its functional significance within the cellular context.

To our best knowledge, this work represents the first comprehensive effort to investigate the proteins derived from non-coding RNA. Several non-coding RNAs like siRNA, piRNA, lincRNA, have not been included in this study and form the future research space.

Though interesting evolutionary and application were derived from this study, it would be interesting to perform experimental validation and generate new biological insights.

## Author Contributions

PKD conceived the idea of making a new class of non-coding proteins from ncRNA molecules. Further, he supervised the work and wrote the final version of the manuscript. SN performed all the computational work and drafted the first version of the manuscript.

## Acknowledgment

SN expresses heartfelt gratitude to UGC-CSIR for the generous provision of the NET-JRF Scholarship and Mr. Manoj Kumar Das for helping in the computational work. PKD expresses warm appreciation to the JNU Administration for their valuable support.

## Notes

### Competing Interest Statement

The authors have declared no competing interest.

